# A suitable murine model for studying respiratory coronavirus infection and therapeutic countermeasures in BSL-2 laboratories

**DOI:** 10.1101/2021.05.28.446200

**Authors:** Ana Cláudia dos Santos Pereira Andrade, Gabriel Henrique Campolina-Silva, Celso Martins Queiroz-Junior, Leonardo Camilo de Oliveira, Larisse de Souza Barbosa Lacerda, Jordane Clarisse Pimenta, Filipe Resende Oliveira de Souza, Ian de Meira Chaves, Ingredy Beatriz Passos, Danielle Cunha Teixeira, Paloma Graziele Bittencourt-Silva, Priscila Aparecida Costa Valadão, Leonardo Rossi-Oliveira, Maisa Mota Antunes, André Felipe Almeida Figueiredo, Natália Teixeira Wnuk, Jairo R. Temerozo, André Costa Ferreira, Allysson Cramer, Cleida Aparecida Oliveira, Ricardo Durães-Carvalho, Clarice Weis Arns, Pedro Pires Goulart Guimarães, Guilherme Mattos Jardim Costa, Gustavo Batista de Menezes, Cristina Guatimosim, Glauber Santos Ferreira da Silva, Thiago Moreno L. Souza, Breno Rocha Barrioni, Marivalda de Magalhães Pereira, Lirlândia Pires de Sousa, Mauro Martins Teixeira, Vivian Vasconcelos Costa

**Affiliations:** Department of Morphology, Institute of Biological Sciences, Universidade Federal de Minas Gerais, Belo Horizonte, MG, Brazil; Department of Biochemistry and Immunology, Institute of Biological Sciences, Universidade Federal de Minas Gerais, Belo Horizonte, MG, Brazil; Department of Microbiology, Institute of Biological Sciences, Universidade Federal de Minas Gerais, Belo Horizonte, MG, Brazil; Department of Physiology and Biophysics, Institute of Biological Sciences, Universidade Federal de Minas Gerais, Belo Horizonte, MG, Brazil; Laboratory on Thymus Research, Oswaldo Cruz Institute, Fiocruz, Rio de Janeiro, RJ, Brazil; National Institute for Science and Technology on Neuroimmunomodulation, Oswaldo Cruz Foundation (Fiocruz), Rio de Janeiro, RJ, Brazil; National Institute for Science and Technology on Innovation on Diseases of Neglected Populations (INCT/IDNP), Center for Technological Development in Health (CDTS), Oswaldo Cruz Foundation (Fiocruz), Rio de Janeiro, RJ, Brazil; Immunopharmacology Laboratory, Oswaldo Cruz Foundation (Fiocruz), Rio de Janeiro, RJ, Brazil; Laboratório de Pesquisas Pré-clínicas, Universidade Iguaçu (UNIG), Rio de Janeiro, RJ, Brazil; Laboratory of Virology, Universidade Estadual de Campinas (UNICAMP), Campinas, SP, Brazil; Department of Metallurgical Engineering and Materials, Federal University of Minas Gerais, School of Engineering, Belo Horizonte, Brazil; Department of Clinical and Toxicological Analysis, Faculty of Pharmacy, Universidade Federal de Minas Gerais, Belo Horizonte, MG, Brazil

## Abstract

Several animal models are being used to explore important features of COVID-19, nevertheless none of them recapitulates all aspects of the disease in humans. The continuous refinement and development of other options of *in vivo* models are opportune, especially ones that are carried out at BSL-2 (Biosafety Level 2) laboratories. In this study, we investigated the suitability of the intranasal infection with the murine betacoronavirus MHV-3 to recapitulate multiple aspects of the pathogenesis of COVID-19 in C57BL/6J mice. We demonstrate that MHV-3 replicated in lungs 1 day after inoculation and triggered respiratory inflammation and dysfunction. This MHV-model of infection was further applied to highlight the critical role of TNF in cytokine-mediated coronavirus pathogenesis. Blocking TNF signaling by pharmacological and genetic strategies greatly increased the survival time and reduces lung injury of MHV-3-infected mice. *In vitro* studies showed that TNF blockage decreased SARS-CoV-2 replication in human epithelial lung cells and resulted in the lower release of IL-6 and IL-8 cytokines beyond TNF itself. Taken together, our results demonstrate that this model of MHV infection in mice is a useful BSL-2 screening platform for evaluating pathogenesis for human coronaviruses infections, such as COVID-19.

## Introduction

*Betacoronavirus* genus belongs to the *Coronaviridae* family and encompasses positive-stranded enveloped RNA viruses. This viral group is broadly distributed among humans and other mammals including rodents, bats, pigs and ruminants (1–3). The high prevalence of betacoronaviruses all over the world, combined with the great genetic diversity and the increased human occupation of isolated ecosystems makes the periodic emergence of novel coronaviruses in zoonotic outbreaks highly probable, after host escape events (1). In fact, over the last two decades, there were three isolated zoonotic outbreaks of severe coronavirus infection in humans including the currently circulating severe acute respiratory syndrome coronavirus 2 (SARS-CoV-2), which is responsible for the ongoing Coronavirus Disease 2019 (COVID-19) pandemic (4). Due to its epidemiological importance, a deeper understanding of viral biology, pathogenesis and the discovery of therapeutic options against betacoronaviruses is imperative.

COVID-19 has a broad spectrum of clinical manifestations. Whereas in most cases SARS-CoV-2 infection is silent or trigger only mild respiratory symptoms, in others it leads to severe conditions including acute respiratory distress syndrome (ARDS), multi-organ failure, systemic inflammation, and death (5,6). We have previously suggested that the development of anti-inflammatory drugs may be useful as co-adjuvant treatment for infectious diseases (7). Indeed, this has gained special interest in the context of COVID-19, especially because the use of steroids have been proven beneficial in hospitalized COVID-19 patients (8). Knowledge of mediators of inflammation contributing to disease initially comes from association studies in humans. Nevertheless, studies in animal models are crucial to define the role of a certain mediator in the pathogenesis of an infection.

Several animal models have been proposed as preclinical platforms to study the pathogenesis of SARS-CoV-2, including mice, ferrets, Syrian hamster and, primates. (9–14). The inherent resistance of wild-type mice to SARS-CoV-2 infection led to the establishment of strategies to adapt these rodents to infection, either by modifying the host or the virus (11,15,16). One of the most common strategies was the infection of transgenic mice expressing human ACE2 (hACE2) using SARS-CoV-2 strains. However, these animal models in general fail to reproduce the severe manifestations of COVID-19 and display a high rate of SARS-CoV-2 replication in non-target tissues (17,18). However, these animal models in general fail to reproduce the severe manifestation of SARS-CoV-2 infection. Due to interspecies differences, a single animal model of COVID-19 is unlikely to reproduce all the vast disease aspects observed in humans. A set of diverse models is are necessary for broader understanding of viral tropism, replication, and dissemination, clinical signs, pathogenesis, and immune response caused by SARS-CoV-2 (19).

The use of other betacoronaviruses, such as the murine coronavirus (MuV), has been suggested as an efficient strategy to emulate many of the key aspects of human CoV (HuCoV) infection biology (20,21). Among MuV, the group previously called murine hepatitis coronavirus (MHV) is the prototype of this genus and a natural pathogen of the *Mus musculus* species. MHV is capable of inducing a severe and lethal disease in mice and some of its variants may have an initial pulmonary tropism after intranasal inoculation, before systemic dissemination. This lung tropism is capable of inducing transient pneumonia that mimics some aspects of the pathogenesis of severe acute respiratory syndrome (20,21). An important advantage of using this murine model is the requirement for biosafety level 2 (BSL-2), which makes this model significantly less costly and safer for screening compounds of therapeutic interest.

The MuV strain MHV-3 has classically been described as a causative agent of severe hepatitis in several strains of mice (22). However, the in-depth characterization carried out in the present study suggests that intranasal infection of wild-type mice by MHV-3 is capable of inducing a transient “SARS-like” disease. We suggest that this animal model may be useful for understanding the pathogenesis of human coronavirus infection and show a critical role for TNF in mediating coronavirus pathogenesis.

## Results

### Intranasally delivered MHV-3 favors pulmonary viral replication in C57BL/6J mice

To characterize a mouse model of coronavirus-induced acute respiratory infection inside a BSL-2 facility, 6- to 7-week-old C57BL/6J mice were inoculated via the intranasal route with 10^3^ plaque-forming units (PFU) of MHV-3 and daily monitored for signs of illness (Fig. 1a). Following infection, mice underwent progressive weight loss starting on the 3^rd^ day post-infection (dpi) and became moribund and died by 6 dpi (Fig. 1b, c). Male and female animals succumbed similarly to the disease as their median survival time upon MHV-3 challenge was 5.5 and 6 days, respectively. MHV-3 infection was also lethal at a lower dose (10^2^ PFU); however, there was a modest delay in weight loss and death compared to higher inoculum (S1 Fig).

**Fig 1:**
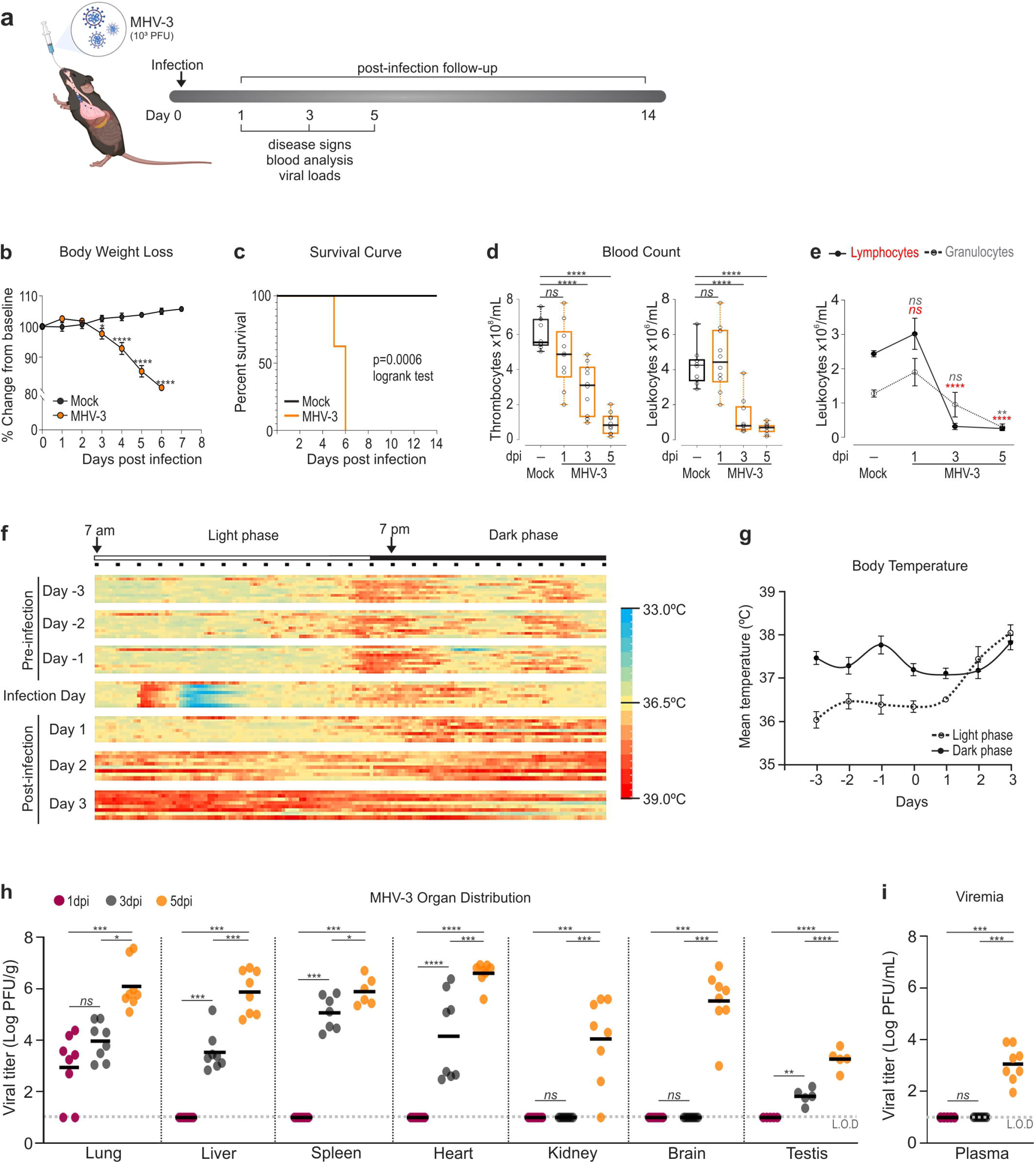
Intranasally inoculated MHV-3 triggers lethal disease in C57BL/6J mice. **a** Experimental design. **b** Body weight change upon infection assessed by two-way repeated measures ANOVA plus Sidak’s multiple comparisons test (mean + S.E.M; n=8). **c** Kaplan-Meier survival curve of infected mice vs. mock controls (n=8). **d** Changes in the number of circulating thrombocytes and leukocytes over 5 days post inoculation (dpi) and represented as box plots. The whiskers go from each the first and third quartile to the minimum or maximum value. Data from each infection time were compared with the mock group by one-way ANOVA plus Dunnett’s multiple comparisons test (n=10). **e** Differential blood count highlighting the sharp infection-related drop of lymphocyte counts that likely guided leukopenia noticed from 3 dpi onwards. Differences between infection groups and mock control were assessed by one-way ANOVA plus Dunnett’s multiple comparisons test (mean + S.E.M; n=6). **f** Heatmap showing the body temperature recorded in a group of 10 mice over 3 days pre- and pos-infection. Note an increase in the mean temperature on dpi 2 and 3 especially in the light phase. **g** Mean + S.E.M of body temperature depicted in (f) according to the time of the day. **h, i** Viral load determined in organ extracts and plasma of MHV-3-infected mice by plaque assay. The results are presented in log10 plaque forming-units (PFU) per gram of tissue or milliliter of plasma. Differences among groups were assessed by Kruskal-Wallis plus Dunn’s post hoc test (n= 5-8). L.O.D= limit of detection. ns= not significant (*P* > 0.05). \**P* < 0.05; \*\**P* < 0.01; \*\*\**P* < 0.001; \*\*\*\**P* < 0.0001.

Intranasal infection of mice with 10^3^ PFU led to progressive thrombocytopenia (Fig. 1d). The number of circulating leukocytes also decreased as the infection progressed, which was mostly driven by the sharp lymphopenia observed from 3 dpi onwards (Fig. 1d, e). Thermal response to viral infection was also observed. As depicted in Fig. 1f, body temperature began to rise slowly 24 hours after virus inoculation. At 2 and 3 dpi, mice reached 37.4 ± 0.3 °C and 38.1 ± 0.2 °C respectively, which was significantly higher than the average temperature recorded over three days before infection (36.3 ± 0.1 °C, p<0.05) (Fig. 1f, g). It is worth noting that major changes in body temperature occurred during the light phase, which is the resting period for nocturnal habit animals (Fig. 1g).

To ascertain whether intranasal delivered MHV-3 favors viral replication in the respiratory system, viral load was assessed in the lung at 1, 3, and 5 dpi and compared with that found in the plasma, liver, spleen, heart, kidney, brain, and testis. Results confirmed the lungs as the initial replication site of MHV-3, as assessed by the presence of infectious virus in this tissue, with viral loads significantly increasing from 1 dpi to 5 dpi (Fig. 1h). Conversely, viruses were recovered from 3 dpi onwards in other organs, with viremia being more stably detected at 5 dpi (Fig. 1h, i). Together, these data show that C57BL/6J mice are highly susceptible to intranasal inoculation of MHV-3 and that lungs are likely the primary site of viral infection and replication.

### MHV-3 triggers inflammation-associated lung damage and respiratory dysfunction

We next investigated whether MHV-3 infection would affect lung morphology and trigger inflammation-associated tissue damage (Fig. 2a). Compared to mock controls, a higher number of cells stained for the pan-leukocyte marker CD45 was found in the lung sections of infected mice after 1 day and especially 3 days after inoculation (Fig. 2b). Flow cytometry results concurred with the recruitment of leukocytes to the lung tissue early during infection, showing a significant rise in the percentage of neutrophils and macrophages/monocytes. On the other hand, we found that the percentage of both CD4^+^ and CD8^+^ T cells in the lung gradually decreased as the infection progressed (Fig. 2 c; S2 Fig). The histopathologic examination of lung sections revealed that MHV-3 infection triggered transient inflammation-associated lung injury. As such, approximately 50% of mice had discrete inflammatory cell infiltration in close association with areas of alveolar edema and hyperemic vessels at 1 day after MHV-3 infection (Fig. 2 d-f). At 3 dpi, the inflammatory infiltrate became robust and widespread throughout the lung tissue, leading to greater histopathological changes. Inflammation foci were frequently seen in the vicinity of bronchioles, in perivascular areas, and in the lumen of hyperemic vessels (Fig. 2 b, d-f). Some blood vessels presented eosinophilic fibrillar material adherent to the vascular wall, which is characteristic of fibrin thrombi. In some samples, areas of necrosis and hemorrhage were also observed (Fig. 2 d). Conversely, the inflammation-associated lung damage in the mouse lung at 5 dpi was milder and encompassed the changes herein portrayed for day 1 after infection (Fig. 2 d-f), suggesting that inflammation was resolving.

**Fig 2:**
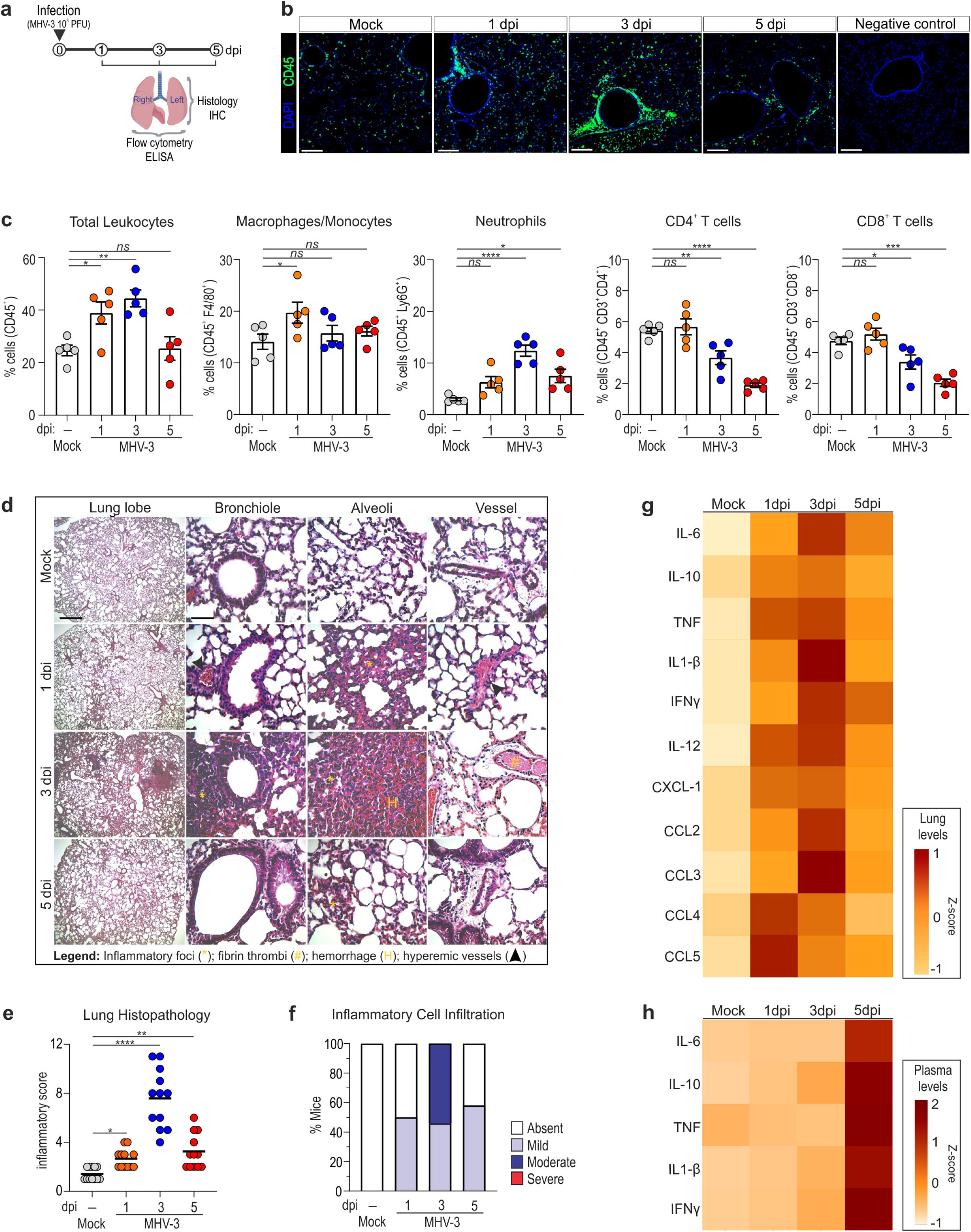
MHV-3 infection triggers inflammation-associated lung injury. **a** Experimental design. **b** Representative confocal 3D image showing high abundance of CD45^+^ leukocytes (green staining) in the lung of mice at 1 and 3 dpi. **c** Flow cytometry analyses showing the percentage of total leukocytes, macrophages/monocytes, neutrophils, and CD4^+^ and CD8^+^ T cells among groups. The percentage of each cell type at 1, 3, and 5 dpi was compared with that found in the mock group by one-way ANOVA plus Dunnett’s multiple comparisons test (mean + S.E.M; n=5). **d** H&E staining of lung sections showing notorious signs of inflammatory injury in infected mice. Scale bars= 50μm. **e** histopathological assessment in relation to overall inflammatory score. Comparisons between mock and infection groups were carried out by Kruskal-Wallis plus Dunn’s post hoc test (n=12). **f** Percent of mice according to the grade of inflammatory cell infiltration. **g** Heatmap evidencing changes in the levels of cytokines and chemokines measured by ELISA in the lung of mock controls and MHV-3-infected mice (n=5). **h** Heatmap showing greatly higher levels of IL-6, IL-10, TNF, IL-1β and IFNγ in the plasma of infected mice at 5dpi (n=5). ns= not significant (*P* > 0.05). \**P* < 0.05; \*\**P* < 0.01; \*\*\**P* < 0.001; *\*\*\*\*P* < 0.0001.

The intrapulmonary concentration of major chemokines (CCL2, CCL3, CCL4, CCL5) and cytokines (TNF, IL-6, IL-1β, IL-12, IFNγ) was markedly increased at 1 dpi and/or 3 dpi and lowered thereafter (Fig. 2 g). In contrast, high levels of the inflammatory mediators (IL-6, IL-10, TNF, IL-1β and IFNγ) were only detected in the blood at 5 dpi (Fig. 2 h). These data suggest that the current infection model exhibits transient pulmonary pneumonia followed by systemic inflammation.

Next, we asked whether these histopathological changes could impact on lung function. By using the whole-body plethysmography method, two control pre-infection measurements of ventilatory parameters were performed in freely moving mice and compared with that recorded on dpi 3 (Fig. 3a). Interestingly, there was an increase in respiratory frequency in MHV-3-infected animals (Fig. 3b) that was characterized by shortened respiratory cycles (expiration/inhalation event; Fig. 3b). In addition, the mechanic of the respiratory system was evaluated invasively in another group of mice at 3 dpi and compared to mock controls (Fig. 3a). There was restricted lung distention after MHV-3 infection, as determined by the lower static compliance of the respiratory system (Fig. 3c). There was also a decrease in vital capacity of the infected group (Fig. 3c). These restrictive aspects of pulmonary function are likely secondary to the pulmonary inflammation and damage. These data suggest that intranasal inoculation of MHV-3 in C57BL/6J mice triggers dysfunction of the respiratory system.

**Figure 3:**
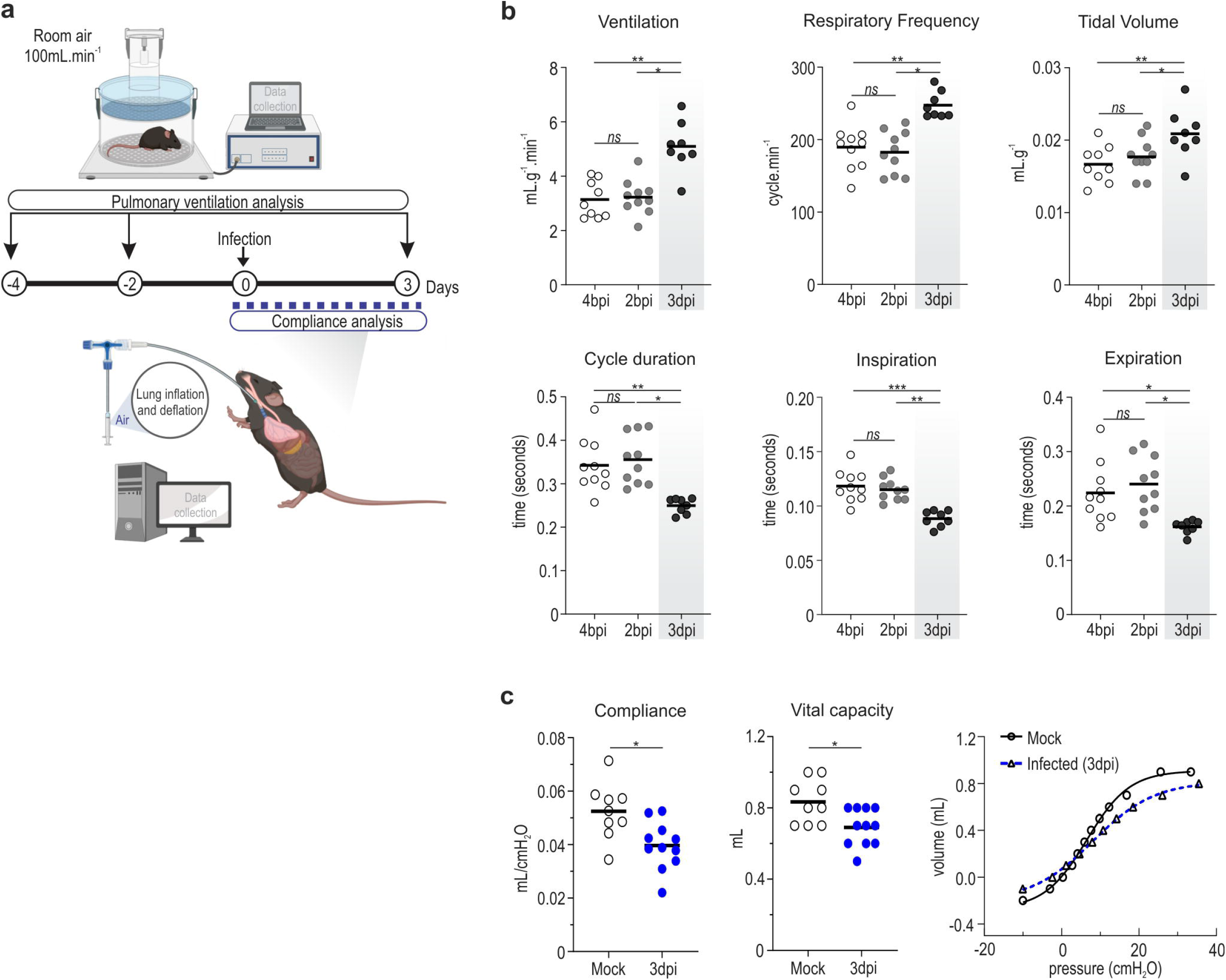
Intranasal infection of MHV-3 impairs lung function. **a** Experimental design. **b** Pairwise comparison of ventilatory parameters recorded 4 and 2 days before infection (bpi) and at 3dpi. Note that mice showed a significant increase in the ventilation along with shortened respiratory cycles upon MHV-3 challenge. One-way repeated measures ANOVA plus Tukey’s post hoc test were applied to assess differences between pre-infection and post-infection periods (n=8). **c** Analysis of pulmonary compliance showing reduced compliance and vital capacity in infected mice compared to mock (unpaired *t* test, n=9-11). Static compliance was calculated from the steepest point of the deflation part of the pressure-volume curve (right graph). ns= not significant (*P* > 0.05). \**P* < 0.05; \*\**P* < 0.01; \*\*\**P* < 0.001; \*\*\*\**P* < 0.0001.

Given the pivotal role of diaphragm contraction for breathing and the recent evidence that SARS-CoV-2 might trigger diaphragm myopathy (23), we next checked the integrity of diaphragm neuromuscular junctions (NMJs) in the current infection model. By combining whole-mount confocal microscopy and computer-assisted image analysis, we showed that NMJs from MHV-3-infected mice at 3 and 5 dpi were significantly smaller and more fragmented at both pre- and post-synaptic endings, in comparison with mock controls (Fig. 4). Although this deterioration of the diaphragm synaptic apparatus may not be sufficient to disrupt pulmonary ventilation in infected mice, its long-term contribution to respiratory muscle weakness needs further attention.

**Figure 4:**
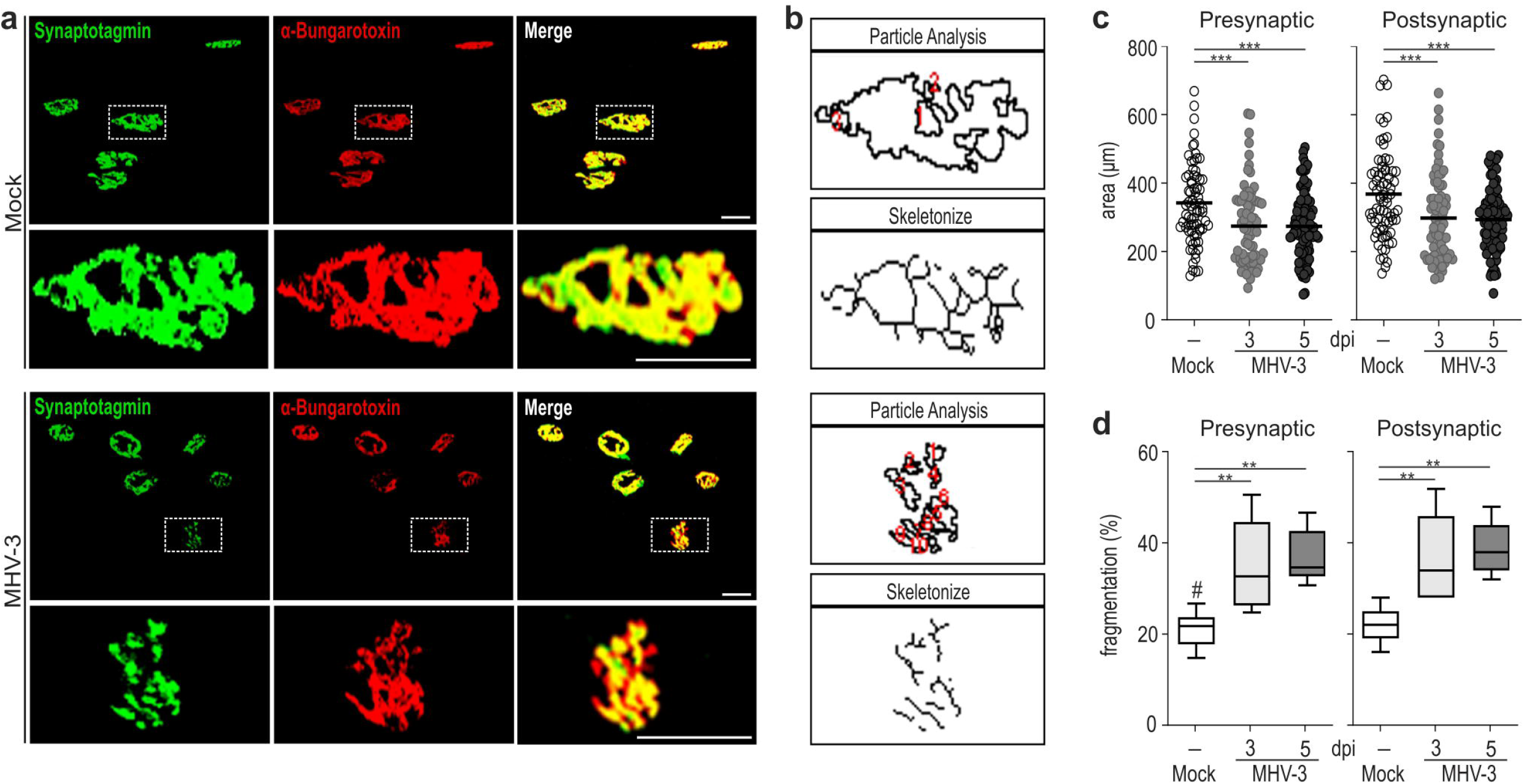
NMJs from diaphragm muscle are smaller and more fragmented in MHV-3-infected mice. **a** 3D confocal images from whole mount diaphragm showing presynaptic (green) and postsynaptic terminals (red) at neuro-muscular junctions (NMJs). Scale bar: 50 μm. **b** Representative images from particle analysis and skeletonization of the NMJ highlighted in figure (a) by dotted squares. **c** Quantification of the area occupied by presynaptic and postsynaptic terminals. 125 NJMs per group were considered in this analysis and the values from infected animals were compared with mock controls by one one-way ANOVA plus Dunnett’s multiple comparisons test. **d** Quantification of the fragmentation percentage found in presynaptic as postsynaptic endings identified by particle analysis. Assessed by one-way ANOVA plus Dunnett’s multiple comparisons test (n=5). \**P* < 0.05; \*\**P* < 0.01; \*\*\**P* < 0.001; \*\*\*\**P* < 0.0001.

### MHV-3 induces extrapulmonary damage

MHV-3 is a murine coronavirus well-known for its hepatotropism (24). Overall, intranasally inoculated MHV-3 triggered hepatitis and liver necrosis, especially at 5 dpi (S3 Fig), the period that preceded systemic clinical manifestations and death of the animals (as shown in Fig. 1 b, c). Consistent with the liver damage, liver function was heavily impaired on dpi 5, as seen by the high serum levels of ALT and the reduced hepatic ability of metabolizing indocyanine dye at this time point (S3 Fig). Other organs such as the brain, small intestine, and colon had minimal to mild leukocyte infiltration at 3 and 5 dpi and were much less affected than the liver (S4 Fig).

Furthermore, in view of recent evidence pointing to the high susceptibility of the testis to both SARS-CoV and SARS-CoV-2 infection (25,26), we evaluated whether MHV-3 could trigger testicular damage. Interestingly, the percentage of altered seminiferous tubules raised significantly after inoculation. The altered sites displayed epithelium sloughing, elevated germ cell apoptosis, and retention of residual bodies (S4 FIg). Also, there appeared to be an increase in the volumetric density of blood vessels and lymph space in the testis interstitium of infected mice (S4 Fig), although such changes did not attain statistical significance. Together, these results indicate that intranasal inoculation of MHV-3 triggers multisystem changes beyond the transient lung damage.

### TNF signaling abrogation significantly reduces lung injury and improves survival in MHV-3-infected mice

Increasing concentrations of TNF have previously been associated with tissue damages triggered by MuCov and human coronaviruses and are currently thought to aggravate COVID-19 severity (6,27,28). Once this cytokine was presently found along with other pro-inflammatory mediators at higher levels within the lung of infected mice (Fig. 2g), we next interrogated the contribution of TNF signaling for the pathogenesis of MHV-3 infection. To this end, we infected mice genetically deficient for TNF receptor type 1 (TNFR1, also known as p55) with 10^3^ PFU of MHV-3 intranasally and compared with wild type (WT) mice. Strikingly, TNFR1 knockout mice (TNFR1 KO) were protected from abrupt weight loss and lethality, with 100% of mice surviving by the end of 14-days follow-up period (Fig. 5 a, b). The progressive leukopenia and thrombocytopenia phenotype observed in WT infected mice were not found in TNFR1 KO (Fig. 5c). Moreover, compared to WT, the TNFR1 KO group had significantly lower viral loads (Fig. 5d), inflammation and injury in their lungs (Fig. 5 e-g).

**Fig 5:**
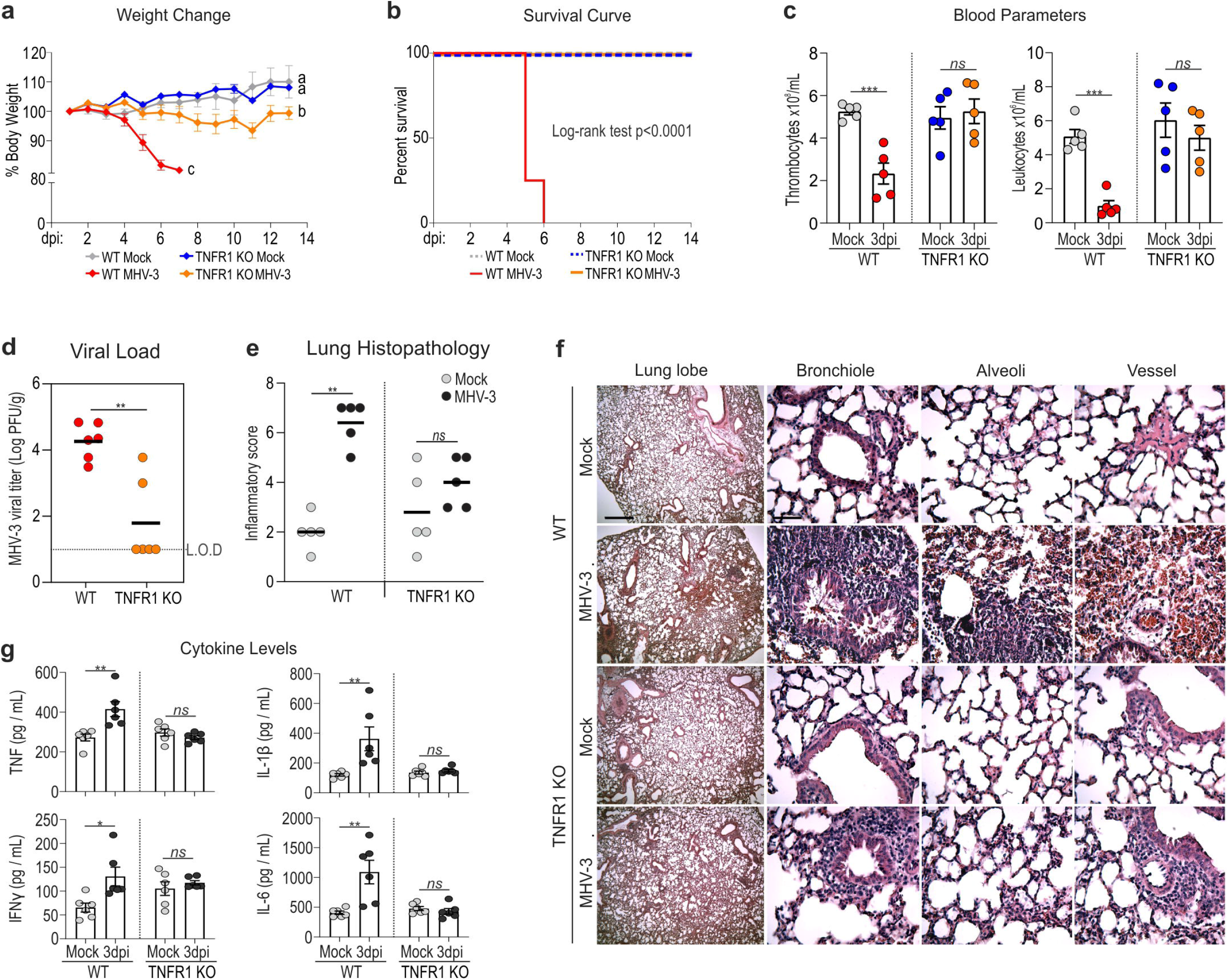
Genetic depletion of TNFR1 prevents MHV-3-mediated lung damage and death in mice. **a** Profile of body weight change in WT and TNFR1 KO mice following 14 days of MHV-3 inoculation. Different letters represent statistically significant differences (p<0.05) among groups assessed by two-way repeated measures ANOVA plus Sidak’s multiple comparisons test (mean + S.E.M; n=6-8). **b** Kaplan-Meier survival curve of WT and TNFR1 KO mice upon infection (n=6-8). **c** Blood analysis showing that the MHV-mediated thrombocytopenia and leukopenia in WT does not occur in TNFR1 KO mice. Mean + S.E.M assessed by unpaired *t* test (n=5). **d** Comparison of the viral load determined by plaque assay in WT vs. TNFR1 KO mice at 3 days after infection. The data were presented in log10 plaque forming-units (PFU) per gram of tissue and assessed by Mann-Whitney test (n=6). L.O.D= limit of detection. **e, f** Comparative histopathology of lungs from mock and infection group (at 3dpi) in relation to the mouse genotype. Statistical comparison among groups were made using the Mann-Whitney test (n=5). **g** Intrapulmonary concentration of TNF, IFNγ, IL-6 and IL-1β cytokines determined ELISA. Note that TNFR1 KO mice were prevented from the MHV-related overproduction of pro-inflammatory cytokines. Comparison between mock vs. infected mice were assessed by unpaired *t* test (mean + S.E.M; n=5-6). ns= not significant (*P* > 0.05). \**P* < 0.05; \*\**P* < 0.01; \*\*\**P* < 0.001; \*\*\*\**P* < 0.0001.

To verify whether such a protective profile against MHV-3 could also be achieved using pharmacological approaches, WT mice were first infected with 103 PFU intranasally and then treated with etanercept, a selective TNF inhibitor. The Etanercept treatment started after 24 hours of infection and was given via two routes: (i) intranasal; or (ii) intraperitoneal (Fig 6 a). Regardless the treatment route, etanercept did not prevent but delayed the body weight loss (Fig. 6b). Similar findings were seen when survival rate was compared among groups, with etanercept-treated mice living on average by 3 days longer than the untreated ones (vehicle group) (Fig. 6 c). It is worth noting that both the local and systemic treatment with etanercept were capable of inhibiting MHV-3 replication in the lungs, with infectious viruses no longer detected in the majority of animals (Fig. 6 d). Consistently, the inflammation-associated lung damage and overproduction of pro-inflammatory cytokines triggered by MHV-3 infection were substantially prevented with etanercept treatment, irrespective of the route of administration (Fig. 6 e-g). Taken together, these findings suggest an important contribution of TNF pathway to the pathogenesis of respiratory coronavirus infection and therapeutic countermeasures in mice.

**Fig 6:**
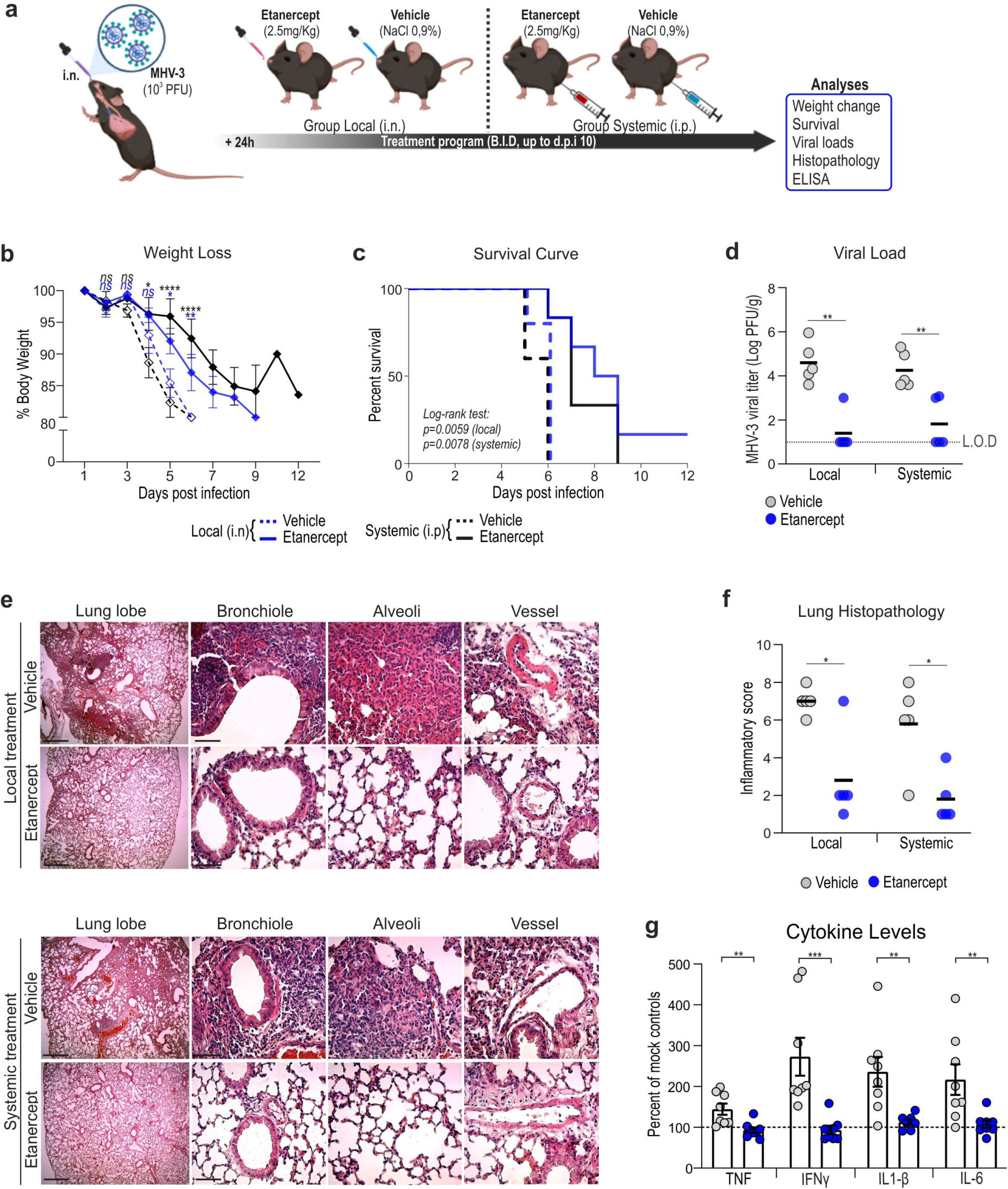
Pharmacologic blockade of TNF cytokine inhibits MHV-3 replication and inflammation-associate injury in the lung of WT mice. **a** Experiment design. Following 24 hours of MHV-3 inoculation, the animals received 2mg/Kg of etanercept or saline (vehicle) twice daily via the intranasal (i.n., local treatment) or intraperitoneal (i.p., systemic treatment) route. Treatment schedule was kept for 10 days in a group of animals to assess body weigh change and lethality, whereas in another group, samples were collected at 3 dpi. **b** Profile of body weight change among groups. Differences between vehicle and etanercept groups were assessed according to the treatment route by two-way repeated measures ANOVA plus Sidak’s multiple comparisons test (mean + S.E.M; n=8). **c** Kaplan-Meier survival analysis of MHV-3-infected mice treated with etanercept vs. vehicle controls (n=8). **d** Viral load determined by plaque assay in the lung of MHV-3-infected mice at 3 dpi treated or not with etanercept Statistical differences were assessed using the Mann-Whitney test (n=5). L.O.D= limit of detection. **e** H&E staining of lung sections showing reduction of inflammation-associated injury signs in etanercept group vs. vehicle controls at 3 dpi. Scale bar= 50μm. **f** Comparative histopathology of lungs according to groups (Mann-Whitney test; n=5). **g** Concentrations of TNF, IFNγ, IL-1β, and IL-6 measured by ELISA assay in the lung of MHV-3-infected mice treated or not with etanercept. The values were normalized to mock group and presented as mean + S.E.M percent of mock controls. Assessed by Mann-Whitney test. n=7-8. ns= not significant (*P* > 0.05). \**P* < 0.05; \*\**P* < 0.01; \*\*\**P* < 0.001; \*\*\*\**P* < 0.0001.

### Blocking TNF decreases SARS-CoV-2 replication and related cellular damage and pro-inflammatory cytokine production in human lung cells

Attempting to translate our findings to huCoV, we next evaluated the effects of etanercept treatment upon SARS-CoV-2 infection *in vitro*. For this, the human epithelial lung cell line Calu-3 was infected with SARS-CoV-2 and then treated with etanercept at different concentrations (0.5, 1, 5, and 10 ng/mL). The results showed that etanercept treatment reduced the SARS-Cov-2-mediated cellular damage in a dose-dependent manner (Fig. 7 b). Etanercept at 5 and 10 ng/mL had also a slight but statistically inhibitory effect on SARS-CoV-2 (Fig. 7c). Moreover, the overproduction of pro-inflammatory cytokines triggered by SARS-CoV-2 infection was significantly reduced in Calu-3 cells treated with etanercept (Fig. 7d). These results suggest that TNF blocking by pharmacologic approaches might also be beneficial against SARS-CoV-2 infection in human lung epithelial cells.

**Figure 7:**
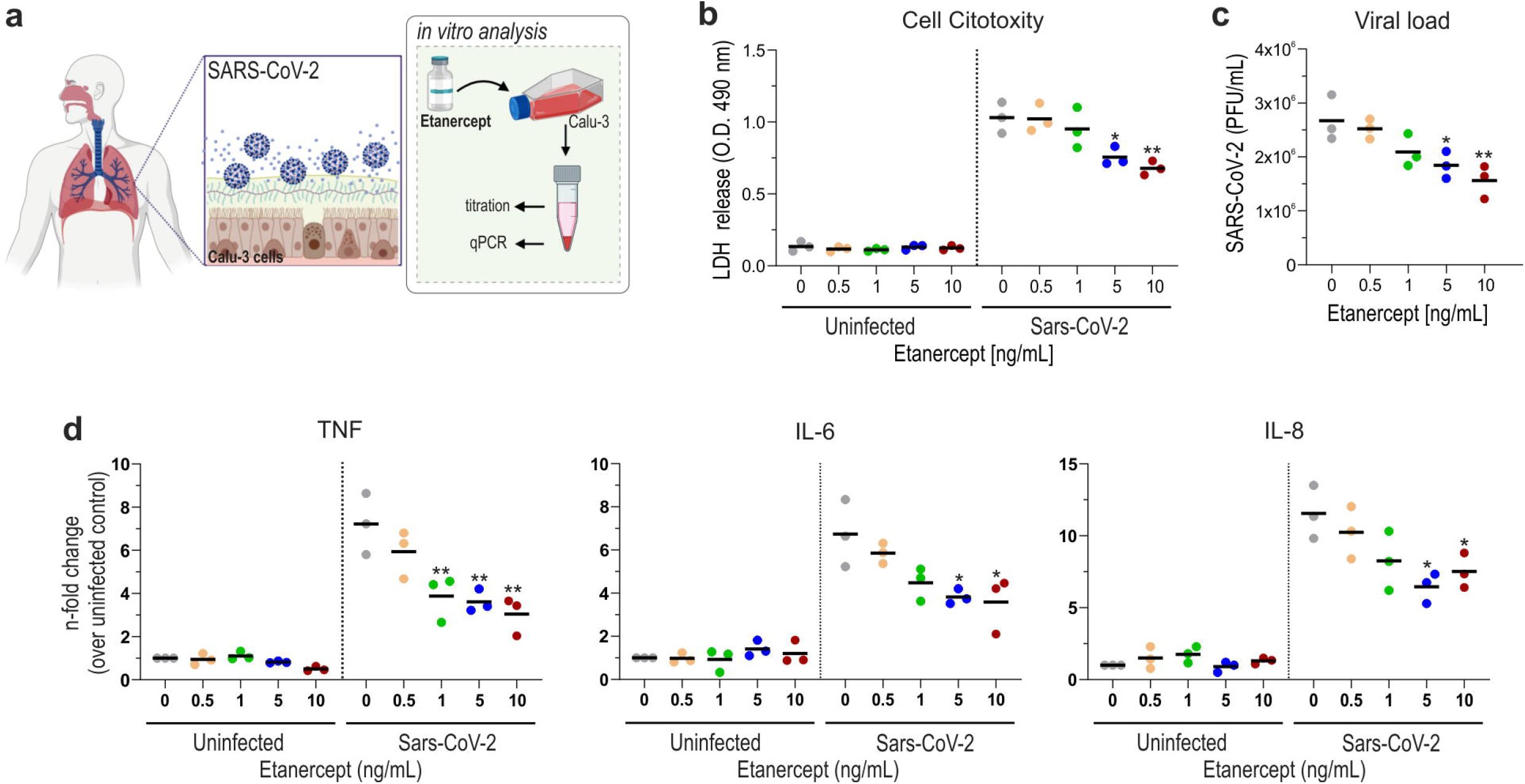
TNF blocking decreases SARS-CoV-2 replication and related cellular damage in human lung epithelial cells. **a** Experiment design. **b** Measurement of cell damage by dosing LDH levels released in cell supernatant of Calu-3 cells treated or not with different concentration of etanercept. **c** Changes in SARS-CoV-2 viral load determined in Calu-3 cells supernatants after treatment with increasing doses of etanercept. **d** ELISA assay showing lower concentration of TNF, IL-6 and IL-8 in CALU-3 infected cells following etanercept treatment. Values from each dose group were compared to untreated controls by One-way ANOVA plus Dunnett’s multiple comparisons test. n= 3 independent experiments. \**P* < 0.05; \*\**P* < 0.01; \*\*\**P* < 0.001; \*\*\*\**P* < 0.0001.

## Discussion

Mouse models have long been used as valuable *in vivo* platforms to investigate the pathogenesis of viral infections and effective countermeasures. The natural resistance of *Mus musculus* to the novel betacoronavirus has launched a race towards the characterization of SARS-CoV-2 infection in other animals (e.g. hamsters, cats, ferrets, bats, and monkeys) as well as the adaptation of the mouse model, by either modifying the host or the virus (19). If on one hand the animal models established to date have advanced knowledge of transmission and host responses to SARS-CoV-2, on the other they require a biosafety level 3 containment and restricted license, which further limit the speed up the development of new therapies. In the present study, we utilized the natural pathogen of mice MHV as a prototype to model betacoronavirus-induced acute lung injure under biosafety level 2 condition. We showed that C57BL/6J mice intranasally inoculated with MHV-3 develop a severe disease mirroring most aspects of severe SARS-CoV-2 infection in patients, which include acute lung damage and respiratory distress preceding systemic inflammation and death. Accordingly, the proposed animal model may provide a useful tool for studies regarding betacoronavirus respiratory infection and related diseases including COVID-19.

MHV-3 replicated in the mouse lung and triggered a robust inflammatory response in the local. The immune response comprised accumulation of neutrophils and macrophages/monocytes accompanied by augmented tissue concentrations of a range of inflammatory mediators, including the chemokines CCL2, CCL3, CCL4, and CCL5 as well as the cytokines TNF, IFNγ, IL1-β, and IL-6. Although these innate cells and cytokines are vitally important in clearing viral infections, increasing epidemiological and experimental studies on SARS-CoV-2 pathogenesis have indicated that the overt host inflammation likely accounts more for the development of severe respiratory illness and fatalities than the virus replication itself (6,29). In line, lung MHV-3 viral burden doubled from the 1^st^ to the 5^th^ day after intranasal inoculation whereas the major histopathological changes, including alveolar edema, hyperemic vessels, hemorrhage, and fibrin thrombi, were frequently observed in the lungs by 3 dpi, which coincides with the peak of pulmonary inflammation. Following 5 days after infection, the local inflammation was resolving, and major lung damages were no longer observed.

Acute respiratory dysfunction is a well-documented clinical sign associated with life-threatening pneumonia, including in severely ill COVID-19 patients (30,31). Although some betacoronaviruses might trigger transient pneumonia in wild-type mice, including the MHV-1, MHV-A59 and MHV-S (20,21), as well as the mouse-adapted SARS-CoV-2 strains (15,16), their impact on respiratory mechanic has scarcely been evaluated. Here, we showed that the transient pneumonia triggered by MHV-3 infection was sufficient to alter the pulmonary mechanics by increasing the respiratory frequency, ventilation, and compliance, thereby recapitulating some clinical signs of severe respiratory coronavirus infection. Although MHV-3-infected mice also presented fragmentation in diaphragm NMJS, there was no sign of respiratory muscle weakness and therefore the respiratory dysfunction most probably is resulting from the virus-induced acute lung damage. Nonetheless, we do not rule out the possibility that coronavirus-induced denervation at diaphragm NMJs could account for respiratory failure in the long term, which could be of particular interest for COVID-19 patients under mechanical ventilatory support (23).

It is noteworthy that after inducing pulmonary changes, MHV-3 spread to multiple organs and triggered a systemic and lethal disease. Virus dissemination has been a hallmark of MHV experimental infection (24), since the MHV strains use the widely expressed carcinoembryonic antigen-related cell adhesion molecule 1 (CEACAM-1) as the entry receptor (32). Although the host receptor used for virus entry differs between murine and human coronaviruses (32,33), the dissemination of MHV-3 to extrapulmonary sites and the subsequent systemic hyperinflammation with high circulating levels of TNF, IFNγ, IL1-β and IL-6 may provide a substantial platform to understand the mechanisms underlying the systemic inflammatory response syndrome (SIRS) and multiple organ failure seen in severe COVID-19 (16,34).

TNF has an important role in the coordination and development of most overexuberant inflammatory responses (6,27,28) and its inhibition can alleviate acute lung injure caused by severe respiratory syncytial virus and influenza virus (35,36). During betacoronavirus infection, the overt TNF release can act synergically with IFN-γ and triggers robust inflammatory cell death (37,38). Increasing studies in both MHV and SARS-CoV-2 have demonstrated that overt pro-inflammatory release including TNF is mediated by the host TLR2-Myd88 cascade (28,29). Therefore, targeting either TNF or TLR2 blockade may provide substantial protection against the pathogenesis of coronavirus infection (29,37). Consistently, we showed that TNFR1 KO mice were fully protected from lung tissue injury and death following MHV-3 intranasal infection. Corroborating these findings, treating infected wild-type mice with the TNF inhibitor etanercept mitigated the MHV-3-induced cytokine release and tissue damage. Moreover, we have investigated whether this cytokine was also relevant during infection with SARS-CoV-2 in human lung cells. Etanercept treatment significantly decreased the cellular damage and pro-inflammatory cytokine release triggered by SARS-CoV-2 infection, reinforcing the evidence that TNF blocking strategy may provide protection against the pathogenesis of betacoronavirus infection.

In summary, the mouse model presented in this work mimics several characteristics of COVID-19 in humans, including efficient virus replication in the lungs, induction of a robust inflammatory response with the recruitment of leukocytes and upregulation of inflammatory mediators. The latter changes were accompanied by a decrease in lung compliance, increase in respiratory frequency and ventilation. In addition, the model made it possible to reinforce the relevant role of TNF for coronaviruses pathogenesis. As well as other animal models already proposed for COVID-19 study, the MHV-3 model also has its limitations. One of them is the difference in the host cell receptor used for viral entry. MHV-3 uses the CEACAM-1 receptor (32) whereas SARS-CoV-2 uses ACE2 (33), which may preclude studies of viral entry or drugs that act on this stage of the replication cycle. Another important limitation is the strong viral tropism towards liver cells and the impairment of the function of this organ in late time of infection. However, our results demonstrating significant lung damage and impairment of pulmonary functions following the first days after infection make our model quite useful to study coronavirus-associated lung inflammation and injury.

## Conclusion

Our findings suggest that MHV-3 is a suitable pathogen to model coronavirus-induced pneumonia in mice and highlight that targeting TNF blockage partially prevents lung damage and disease phenotype upon MHV-3 infection. In addition, this model features a lower cost and safer pre-clinical *in vivo* tool capable to recapitulate many aspects of human coronavirus diseases, such as COVID-19.

## Materials and Methods

### Cells, viruses and plaque assay

Vero E6 (ATCC^®^ CRL-1586), L929 (ATCC^®^ CCL-1), and Calu-3 (ATCC^®^ HTB-55) cells were cultured under a controlled atmosphere (37^a^C, 5% CO_2_) in high glucose DMEM (Vero and L929) or MEM (Calu-3) supplemented with 7% fetal bovine serum (FBS), 100 U/mL penicillin and 100 μg/mL streptomycin. MHV-3 strain was kindly provided and sequenced (GenBank accession no. MW620427.1, Ref.(39) by Dr. Clarice Arns and Dr. Ricardo Durães-Carvalho from the Universidade Estadual de Campinas (UNICAMP, Brazil), and propagated in L929 cells. SARS-CoV-2 was expanded in Vero E6 cells from an isolate contained on a nasopharyngeal swab obtained from a confirmed case of COVID-19 in Rio de Janeiro, Brazil (GenBank accession no. MT710714), accordingly to WHO guidelines. For viral titration, 100 μL of serially diluted virus suspension, plasma samples and tissue homogenates (1:9; tissue: DMEM) were inoculated onto a confluent monolayer of L929 cells grown in 24 well-plates (for MHV-3) or Vero E6 cells (for SARS-CoV-2). After gentle shaking for 1 h (4x 15 min), samples were removed and replaced with the overlay medium (DMEM containing 1,6% carboxymethylcellulose, 2% FBS, and 1% penicillin-streptomycin-glutamine) and kept for 2 days (for MHV-3) or 3 days (for SARS-CoV-2), at 37°C and 5% CO_2_. Then, cells were fixed with 10% neutral buffered formalin for 1h and stained with 0.1% crystal violet. Virus titers were determined as plaque-forming units (PFU).

### Mouse strains

The animal’s experimental procedures were carried out with mixed groups (males and females) of mice aged 6 to 7 weeks and received the approval of the Ethical Committee for Animal Experimentation of the Universidade Federal de Minas Gerais (UFMG) (process No. 190/2020). Wild-type C57BL/6 (Central Animal House of the UFMG) and TNF receptor Knockout mice (TNFRp55^-/-^, Jackson Laboratories, stock No. 002818) were housed in individually ventilated cages placed in an animal care facility at 24 ± 2 °C on a 12-h light/ 12-h dark cycle, receiving *ad libitum* access to water and food.

### MHV-3 infection

Mice were lightly anesthetized with an intraperitoneal injection of ketamine (50 mg/kg): xylazine (5 mg/kg) and received an intranasal inoculation of 30 μl sterile saline solution loaded or not (Mock controls) with MHV-3 at different concentrations (3×10^1^ to 3×10^4^ PFU). Signs of disease, including ruffled fur, back-arching, weight loss, facial edema and lack of activity were monitored daily for up to 14 days post inoculation (dpi).

### Tissue collection

Following anesthesia (ketamine: xylazine, 80 mg/kg: 10 mg/kg, i.p), mock controls and infected mice at 1, 3, and 5 dpi were euthanized by cervical dislocation and blood samples were collected from the abdominal vena cava and placed in EDTA-coated tubes. Then, lungs were harvested, briefly rinsed in cold PBS (pH 7.4), and the right lobes were snap-frozen in liquid nitrogen. In contrast, the left lobe was fixed by immersion in 10% neutral buffered formalin solution, unless otherwise mentioned. Formalin-fixed and frozen fragments of liver, heart, brain, small intestine, colon, spleen, and testis were also processed for additional analyses.

### Hematological evaluation

The number of circulating thrombocytes and leukocytes was determined in blood samples using the Nihon Kohden’s Celltac MEK-6500K hemocytometer.

### Histopathology

Formalin-fixed and paraffin-embedded (FFPE) tissues were sectioned into 5 μm thickness slices, stained with hematoxylin-eosin, and examined under light microscopy. The inflammation-mediated injury in mouse lungs was determined by a pathologist (C.M.Q.J) blinded to the experiment, by employing a scoring system encompassing: (i) airway inflammation (up to 4 points); (ii) vascular inflammation (up to 4 points); (iii) parenchyma inflammation (up to 5 points); and general neutrophil infiltration (up to 5 points) (40). Histopathological assessments were also performed in FFPE samples of liver, brain, small intestine, and colon following previously established criteria (41–44). For testis analysis, the tissue was fixed by immersion in Bouin’s solution, embedded in glycol methacrylate resin, sectioned at 3 μm thickness, and then stained with toluidine blue prior to stereological and morphometric assessments, as described (45).

### Chemokine, cytokine and LDH dosage

Lung homogenates were acquired by homogenizing 40 mg of frozen tissue in 400 mL of chilled cytokine extraction buffer (100 mM Tris pH 7.4, 150 MM NaCl, 1 mM EGTA, 1mM EDTA, 1% Triton X-100, 0,5% sodium deoxycholate, 1% protease inhibitor cocktail). After centrifugation (14000 g, 15 min, 4°C), the supernatant was collected and submitted along with plasma samples to measure the concentration of TNF (cat No. DY410), IFN-γ (cat No. DY485), IL-10 (cat No. DY417), IL-6 (cat No. DY406), IL-1β (cat No. DY401) and IL-12 p70 (cat No. DY419) by using the mouse DuoSet ELISA system (R&D Systems). In addition, chemokines such as CXCL-1 (cat No. DY473), CCL-2 (cat No. DY479), CCL3 (cat No. DY450), CCL-4 (cat No. DY451), and CCL-5 (cat No. DY478) were also determined by mouse DuoSet ELISA (R&D Systems) in lung homogenates.

Calu-3 supernatant was obtained after 48 hours of SARS-CoV-2 infection (MOI: 0,1) with or without treatment with Etanercept (0,5-10 ng/mL). Cells were treated after viral adsorption. The levels of IL-6 (cat. No. DY206), IL-8 (cat. No. DY208) and TNF-α (cat. No. DY210) were quantified in the supernatants from uninfected and SARS-CoV-2-infected Calu-3 cells by ELISA (R&D Systems), following manufacturer’s instructions, and results are expressed as n-fold relative to uninfected cells. Cell death was determined according to the activity of lactate dehydrogenase (LDH) in the culture supernatants using a CytoTox1 Kit according to the manufacturer’s instructions (Promega, USA).

### Immunofluorescence

Lung tissues were fixed with 4% paraformaldehyde for 24 h, dehydrated in 30% sucrose, covered with OCT compound, frozen in liquid nitrogen-cooled isopentane, and then sectioned at 15 μm thickness. Following permeabilization in PBS / 0,5% Triton X-100, the cryosections were incubated for 1 h in the blocking solution (PBS containing 5% goat serum and 5 μg/mL mouse BD Fc Block™) and then labeled overnight at 4°C with the APC-conjugated rat anti-mouse CD45 antibody (1:100 dilution, BD Pharmingen, cat No. 559864). Cell nucleus was stained with DAPI. Cell nuclei were stained with DAPI (1 μg/mL, Sigma-Aldrich, Cat No. D9542). Fluorescent signals were evaluated by confocal microscopy using an inverted Nikon Eclipse Ti microscope coupled to an A1 scanning head.

### Flow cytometry

Lungs were minced into small pieces and allowed to digest with gentle agitation (80 rpm, 37°C, 45 min) onto canonical tubes containing 5 mL of digestion buffer (0.5 mg/ml collagenase and 20 μg/ml DNAse diluted in RPMI medium). After, the cell suspension was passed through the cell strainer pore size 70-μm (BD Biosciences, San Jose, CA) and the remaining erythrocytes were lysed with ACK buffer (Invitrogen, Carlsbad, CA). 1×10^6^ cells were blocked with mouse BD FC block™ (5 μg/mL, BD Pharmingen, Cat No. 553141) and then stained using fluorescent-labeled monoclonal antibodies as follows: Ly6G-BV421 (1:200, BioLegend, Cat No. 127627); CD45-FITC (1:200, BD Pharmingen, Cat No. 5530801); F4/80-PE-Cyanine7 (1:100, Invitrogen, Cat No. 25-4801-82); CD4-PE (1:100, BioLegend, Cat No. 100408); CD3-PerCP-Cyanine5.5 (1:100, BioLegend, Cat No. 100218); CD8-APC (1:100, Invitrogen, Cat No. 17-0081-82). The acquisition was carried out in a BD FACSCanto II cell analyzer and analyzed using FlowJo software (Tree Star, Ashland, OR).

### Liver function analyses

Serum concentrations of alanine aminotransferase (ALT) and Indocyanine green (ICG) were used to estimate the liver function status, as previously described (46). In addition, liver necrosis was evaluated by intravital microscopy after labeling free DNA. To this end, 20 minutes before of *in vivo* imaging, mice received an intravenous injection of 1 mg sytox orange diluted in 0.1 mL of sterile saline solution (47).

### Measurement of body temperature

Ten days before MHV-3 inoculation, a group of 10 mice underwent surgery to implant a temperature probe (mini dataloggers, SubCue, Calgary, AB, Canada) into the abdominal cavity. Temperature probes were programmed to start data acquisition at 7 a.m. on the fifth day after surgery. Upon starting, animal’s abdominal temperature was monitored every ten minutes (sample rate) throughout the days that comprised pre- and post-infection procedures. On the fourth-day post infection, mice were euthanized, the dataloggers were collected and the body temperature readings of each animal were downloaded and analyzed (SubCue software, Calgary, AB, Canada).

### Pulmonary ventilation

The whole-body plethysmography method (barometric method) was employed to assess pulmonary ventilation (*in vivo*, freely moving animals) at 4 and 2 days before infection (dbi) as well as at 3dpi. Mice were placed in a cylindrical chamber (mice plethysmography chamber; Bonther, Ribeirao Preto, SP, Brazil) flushed with room air (1000 mL·min^-1^) and allowed to move freely and acclimate for at least 30 min. To measure pulmonary ventilation, the chamber was totally closed and air flow was stopped for a short moment (~ 2 min) while the breathing-related pressure oscillations were detected (differential pressure transducer and amplified; MLT141 spirometer, Power Lab, AdInstruments, NSW, Australia) and recorded with a specific software (LabChart v.7, AdInstruments, NSW, Australia). The ventilation was calculated as the product of the tidal volume (mL BTPS·g^-1^, Ref. (48) and respiratory frequency (breath cycles·min^-1^), corrected by the body weight (g). From the ventilatory recording traces, the inspiration and expiration time as well as total respiratory cycle duration were also calculated.

### Respiratory mechanics

Mice were divided into two groups (mock n=9 and infected n=11) and three days post infection animals were deeply anesthetized until respiratory arrest. Mice were tracheostomized and a polyethylene tube (P50) was inserted into the trachea. The pressure-volume curve was made by injecting air volume in a step-wise manner (using the 3 mL glass syringe), with 0.1 mL increments until intratracheal pressure peaked at approximately 35 cmH_2_O. In the deflation limb, the system was deflated in the same volume steps until the pressure reached approximately –15 cmH_2_O and finally inflated again to resting lung volume. Signals were acquired and recorded on the PowerLab software (LabChart v7, AdInstruments, NSW, Australia). The vital capacity was determined by maximal inflation (lung volume at 35 cmH_2_O) and the static compliance of the total respiratory system (expressed as mL/cmH_2_O) were measured at the steepest point of the deflation limb of the pressure-volume curve.

### Neuromuscular junction (NMJs) analysis of diaphragm muscle

Whole-mount diaphragms were used to evaluate changes in NMJs of MHV-3-infected mice and mock controls. Sample processing and labeling of NMJs were performed based on a previous protocol (49). The pre-synaptic and post-synaptic terminals were stained using the monoclonal anti-synaptotagmin antibody (1:250 dilution, Developmental Studies Hybridoma Bank; cat No. 3H2 2D7) and the tetramethylrhodamine-conjugated α-bungarotoxin (1:1000 dilution; Invitrogen, cat No. T1175), respectively. After Z-stack imaging in a Zeiss LSM 880 confocal microscope, 25 NMJs per animal were assessed by particle analysis to estimate the fragmentation index at pre- and post-synaptic endings (50).

### Etanercept treatment

For *in vivo* TNF inhibition (therapeutical etanercept treatment) during MHV-3 infection, mice were divided into four groups (n = 6 per group, 3 males and 3 females) as follows: a) Vehicle group: mice were i.p. inoculated with 30 μl 0.9% NaCl 24 h before and every day, during 10 days, B.I.D, after intranasal infection with 10^3^ PFU/mouse of MHV-3; b) Systemic treated group (therapeutical scheme): mice were i.p. inoculated with 2.5 mg/kg of etanercept (Enbrel^®^ Pfizer) dissolved 30 μl 0.9% NaCl 24 h after and every day, during 10 days (B.I.D) after intranasal infection with 10^3^ PFU/mouse of MHV-3; c) Local treated group (therapeutical scheme): etanercept was administered by intranasal route in the same dosage as described in b; d) Uninfected treated group (Mock-treated) which received 2.5 mg/kg of etanercept in the same manner as described in b; e) Uninfected group (Mock controls): received only the 0.9% NaCl and was not infected. Lungs of the animals were collected for histopathological analysis and viral loads analysis at 3dpi. Mice survival and body-weight loss were also evaluated.

### Statistical analyses

Graph Pad Prism 8.0 software was used for statistical analysis. First, data distribution was assessed by the Shapiro-Wilk test and Q-Q plots. Parametric comparisons between two or more groups were done using Student-*t* test or one-way ANOVA, respectively, or by using Mann-Whitney or Kruskal-Wallis test to assess differences between two or more non-parametric datasets. Survival rates among groups were determined by Kaplan-Meier survival analysis. Finally, the body weight changes triggered by MHV-3 were compared among groups by two-way repeated measures ANOVA. Data were presented as mean + S.E.M. Differences were considered statistically significant when p<0.05.

## Acknowledgements

This work was financially supported by the Grants from Coordenação de Aperfeiçoamento de Pessoal de Nível Superior – CAPES/Brazil (grant n° 88887.507173/2020-00, to M.M.T), Fundação de Amparo à Pesquisa do Estado de São Paulo – FAPESP/Brazil (grant n° 2019/01255-9, to R.D.C), Ferring COVID-19 Investigational Grant (grant n° FIN0042393, to G.M.J.C), and the L’Oréal-Unesco-ABC For Women in Science Program (grant to V.V.C). This work also received support from the National Institute of Science and Technology in Dengue and Host-microorganism Interaction (INCT dengue), sponsored by the Conselho Nacional de Desenvolvimento Científico e Tecnológico (CNPq, Brazil) and the Fundação de Amparo à Pesquisa do Estado de Minas Gerais (FAPEMIG, Brazil). The authors also acknowledge Dr. Leda Quercia Vieira for the donation of the TNFR1 KO mice and Ilma Marçal de Souza for her technical assistance with experiments.

## Author contributions

**Study design:** A.C.S.P.A., G.H.C.S., C.M.Q.J., L.C.O, L.P.S., M.M.T., G.S.F.S., V.V.C. **Performed experiments:** A.C.S.P.A., G.H.C.S., C.M.Q.S., L.C.O, L. S.B.L., J.C.P., F.R.O.S., I.M.C., I.B.P., D.C.T., P.G.B.S., P.A.C.V., L.R.O., M. M.A., A.F.A.F., N.T.W., A.C.F., J.R.T., A.C. **Data analysis:** A.C.S.P.A., G.H.C.S., L.P.S., G.S.F.S., G.M.J.C., C.G., T.M.L.S., R.D.C., V.V.C. **Intellectual, technical and resource support:** M.M.T., C.A.D., C. W. A., P.P.G.G., G.B.M., T.M.L.S., M. M. P., B. R. B. **Wrote the first draft of the paper:** A.C.S.P.A., G.H.C.S., C.M.Q.J., L.C.O, L.P.S., G.S.F.S., C.G., V.V.C. All authors reviewed and approved the final version of the manuscript.

## Supporting information

**S1 Fig: Body weight change and survival rates according to MHV-3 inocula**. **a** Profile of weight loss following MHV-3 inoculation. Differences between mock controls and infection groups were assessed by two-way repeated measures ANOVA plus Sidak’s multiple comparisons test (mean + S.E.M; n=8). **b** Kaplan-Meier survival analysis comparing the percent survival with increasing load of MHV-3 (n=8). ns= not significant (*P* > 0.05). \**P* < 0.05; \*\**P* < 0.01; \*\*\**P* < 0.001; \*\*\*\**P* < 0.0001.

**S2 Fig: Gating strategy for the analysis of leukocyte populations in the mouse lung.** The results from this analysis were depicted in Fig 2. **a** Representative flow cytometry plots showing the gating scheme for isolating total leukocytes (CD45^+^), macrophages/monocytes (CD45^+^F4/80^+^), neutrophils (CD45^+^Ly6G^+^), CD4^+^ (CD45^+^CD3^+^CD4^+^) and CD8^+^ (CD45^+^CD3^+^CD8^+^) T cells from the lung of MHV-3-infected mice and mock controls. **b** Representative flow cytometry histograms of target cell types according to groups.

**S3 Fig: MHV-3 triggers liver dysfunction following 5 days of infection. a** H&E staining of liver sections showing significant signs of tissue damage at 3 pi and especially at 5 dpi. Areas morphologically compatible with necrosis were indicated by dotted circles. #= hemorrhage. Scale bar= 50μm. **b** Z-stack reconstruction showing extracellular DNA deposition in necrotic areas (arrowheads) identified by Sytox staining (in red). Autofluorescence (blue) was used to highlight the liver parenchyma. Scale bar= 300μm. **c** Mean injury scores compared among groups by Kruskal-Wallis plus Dunn’s post hoc test. **d** Percent of mice in each group according to the necrosis severity (n=7-10). **e** Estimation of liver dysfunction analyzed by serum concentration of alanine aminotransferase (ALT) and indocyanine green (ICG) dye. Statistical differences among groups were assessed by one-way ANOVA plus Tukey’s post hoc test (n=5-6). ns= not significant (*P* > 0.05). \**P* < 0.05; \*\**P* < 0.01; \*\*\**P* < 0.001; *\*\*\*\*P* < 0.0001.

**S4 Fig: Extra-pulmonary changes triggered by MHV-3 inoculated via the intranasal route. a** H&E staining of the brain, small intestine and colon sections showing mild signs of inflammatory injury in infected mice. Original magnification, X400. **b** Mean injury scores compared among groups by Kruskal-Wallis plus Dunn’s post hoc test (n=5-6). **c** Toluidine blue staining of testicular cross-sections showing morphological alterations in infection groups. ST= seminiferous tubule; IC= intertubular compartment *= epithelium sloughing; Arrowheads= indicate germ cell apoptosis; Reddish line= blood vessels; Bluish line= lymph space. Scale bar= 50μm. **d-e** Comparative morphometric analysis conducted in the tubular (d) and intratubular (e and f) of mock vs. MHV-infected mice. Assessed by one-way ANOVA plus Tukey’s post hoc test (Mean + S.E.M, n= 5). ns= not significant (*P* > 0.05). \**P* < 0.05; \*\**P* < 0.01; \*\*\**P* < 0.001; \*\*\*\**P* < 0.0001.

