## Supplementary figures and images for "A suitable murine model for studying respiratory coronavirus infection and therapeutic countermeasures in BSL-2 laboratories"

### S1_Fig.jpg

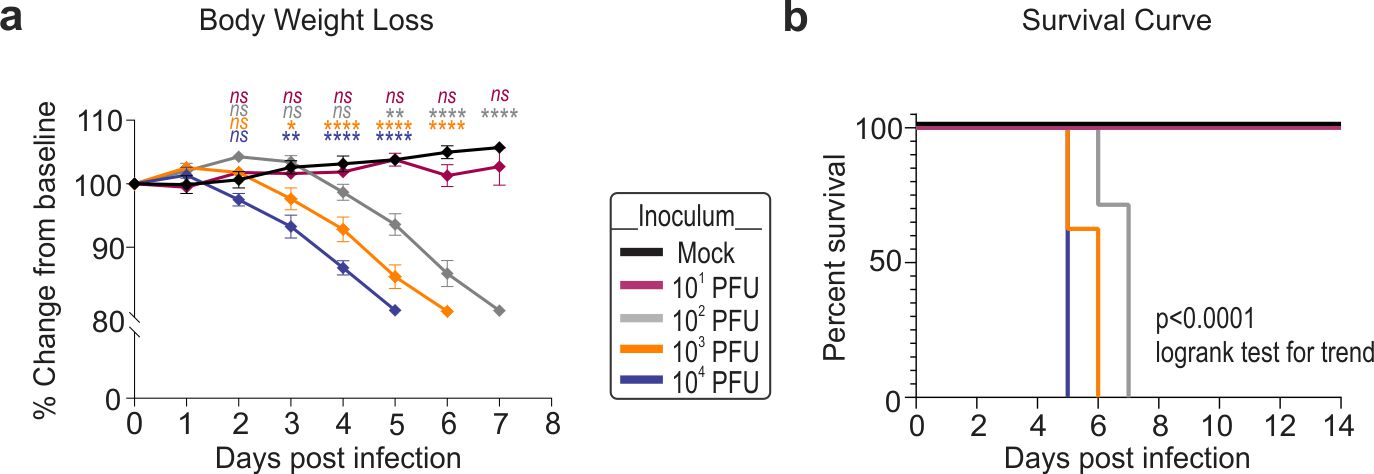

### S2_Fig.jpg

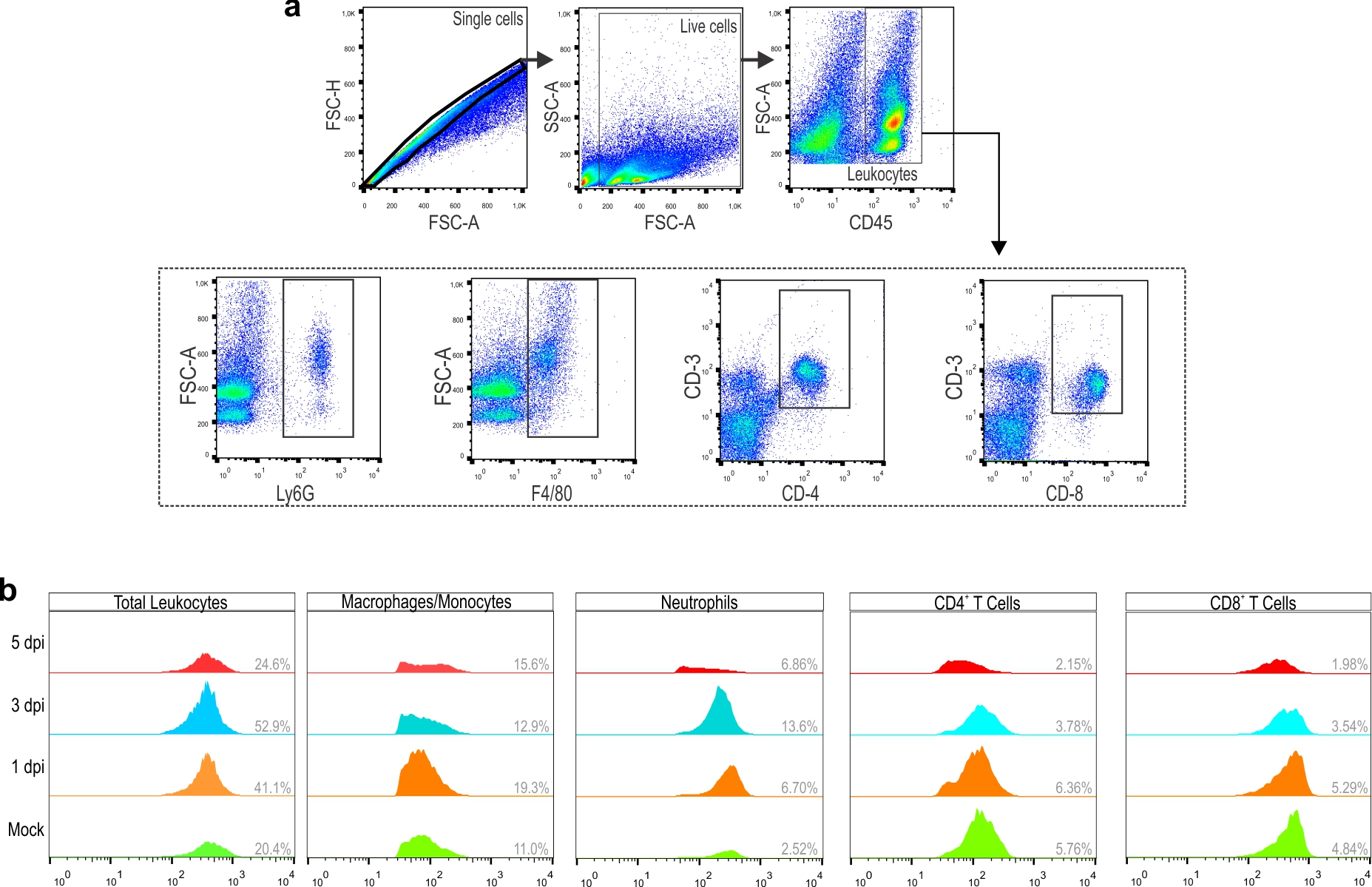

### S3_Fig.jpg

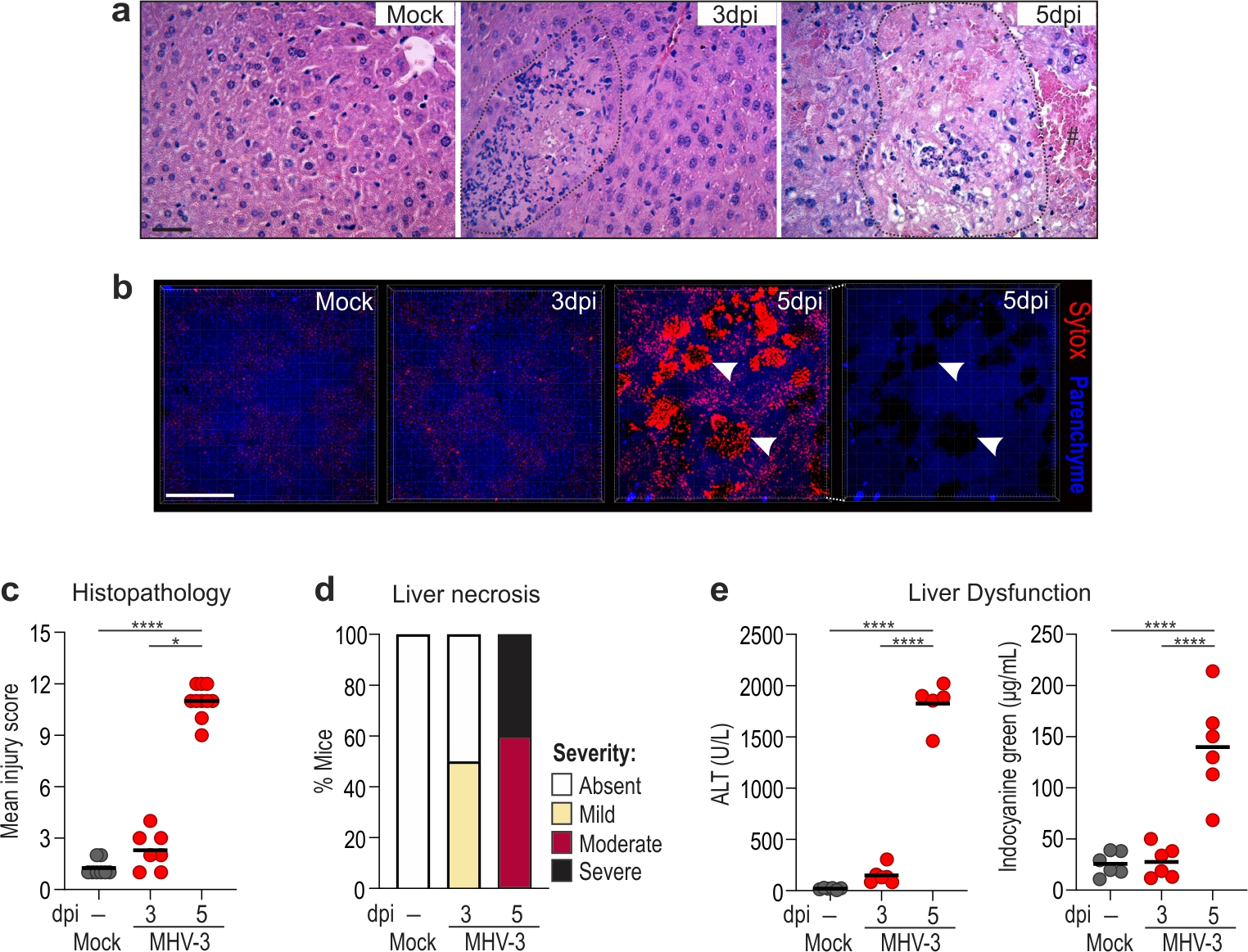

### S4_Fig.jpg

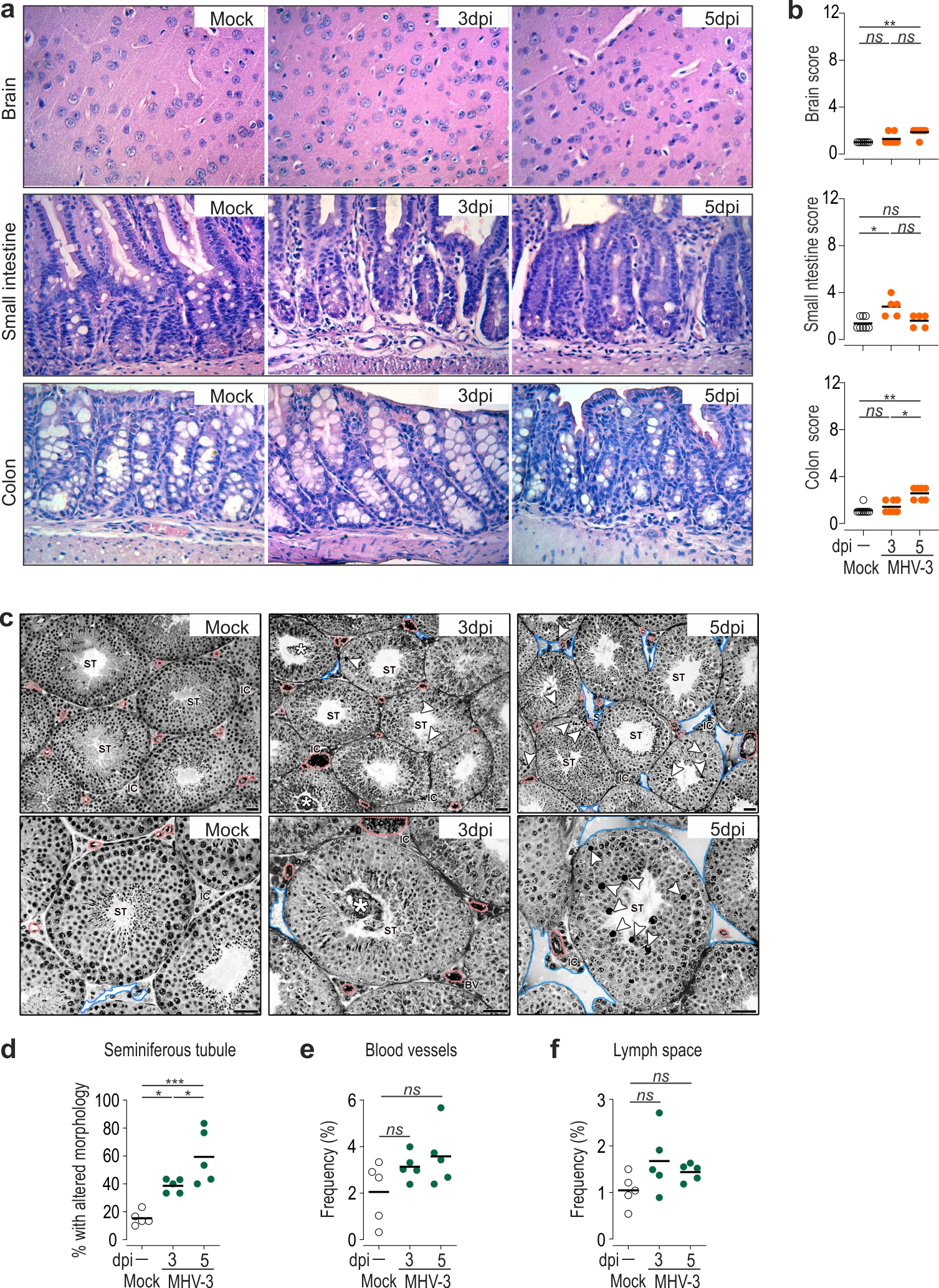
